# A systematic review and evaluation of Zika virus forecasting and prediction research during a public health emergency of international concern

**DOI:** 10.1101/634832

**Authors:** P-Y Kobres, JP Chretien, MA Johansson, J Morgan, P-Y Whung, H Mukundan, SY Del Valle, BM Forshey, TM Quandelacy, M Biggerstaff, C Viboud, S Pollett

## Abstract

**INTRODUCTION:** Epidemic forecasting and prediction tools have the potential to provide actionable information in the midst of emerging epidemics. While numerous predictive studies were published during the 2016-2017 Zika Virus (ZIKV) pandemic, it remains unknown how timely, reproducible and actionable the information produced by these studies was.

**METHODS:** To improve the functional use of mathematical modeling in support of future infectious disease outbreaks, we conducted a systematic review of all ZIKV prediction studies published during the recent ZIKV pandemic using the PRISMA guidelines. Using MEDLINE, EMBASE and grey literature review, we identified studies that forecasted, predicted or simulated ecological or epidemiological phenomenon related to the Zika pandemic that were published as of March 01, 2017. Eligible studies underwent evaluation of objectives, data sources, methods, timeliness, reproducibility, accessibility and clarity by independent reviewers.

**RESULTS:** 2034 studies were identified, of which n = 73 met eligibility criteria. Spatial spread, R_0_ (basic reproductive number) and epidemic dynamics were most commonly predicted, with few studies predicting Guillain-Barré Syndrome burden (4%), sexual transmission risk (4%) and intervention impact (4%). Most studies specifically examined populations in the Americas (52%), with few African-specific studies (4%). Case count (67%), vector (41%) and demographic data (37%) were the most common data sources. Real-time internet data and pathogen genomic information were used in 7% and 0% of studies, respectively, and social science and behavioral data were typically absent in modeling efforts. Deterministic models were favored over stochastic approaches. Forty percent of studies made model data entirely available, 29% provided all relevant model code, 43% presented uncertainty in all predictions and 54% provided sufficient methodological detail allowing complete reproducibility. Fifty-one percent of predictions were published after the epidemic peak in the Americas. While the use of preprints improved the accessibility of ZIKV predictions by a median 119 days sooner than journal publication dates, they were used in only 30% of studies.

**CONCLUSIONS:** Many ZIKV predictions were published during the 2016-2017 pandemic. The accessibility, reproducibility, timeliness, and incorporation of uncertainty in these published predictions varied and indicates that there is substantial room for improvement. To enhance the utility of analytical tools for outbreak response, it is essential to improve the sharing of model data, code, and preprints for future outbreaks, epidemics and pandemics.

**Author summary:** Researchers published many studies which sought to predict and forecast important features of Zika virus (ZIKV) infections and their spread during the 2016-2017 ZIKV pandemic. We conducted a comprehensive review of such ZIKV prediction studies and evaluated their aims, the data sources they used, which methods were used, how timely they were published, and whether they provided sufficient information to be used or reproduced by others. Of the 73 studies evaluated, we found that the accessibility, reproducibility, timeliness, and incorporation of uncertainty in these published predictions varied and indicates that there is substantial room for improvement. We identified that the release of study findings before formal journal publication (‘pre-prints’) increased the timeliness of Zika prediction studies, but note they were infrequently used during this public health emergency. Addressing these areas can improve our understanding of Zika and other outbreaks and ensure that forecasts can inform preparedness and response to future outbreaks, epidemics and pandemics.

## Introduction

Zika virus (ZIKV) is a positive sense RNA flavivirus primarily transmitted through the *Aedes aegypti* mosquito (1–3). While the majority of ZIKV infections are asymptomatic or present as a self-limiting febrile illness, strong evidence links ZIKV infection with microcephaly and a range of other birth defects including limb deformity and retinopathy (4, 5). ZIKV is also associated with Guillian-Barre syndrome, and a spectrum of other neurological disorders including meningoencephalitis and acute myelitis (6–9). ZIKV was discovered in Uganda in a febrile non-human primate in 1947 (10), and the first human case was detected in Nigeria in 1953 (11). ZIKV outbreaks were detected in South East Asia and the Pacific Islands in the early 21^st^ century (12–16) followed by wide spread epidemics in the Americas from late 2014 onward with a cumulative count of 583,144 suspected and 223,336 laboratory-confirmed Zika cases reported across 49 countries and territories by the end of 2017 (17, 18).

The Director-General of the World Health Organization declared the ZIKV pandemic a public health emergency of international concern (PHEIC) on February 1, 2016 (19). The urgency for immediate, coordinated global response was further accelerated by the Olympic and Paralympic games set to take place in Rio De Janeiro, Brazil during August 2016 (20). As public health and medical research efforts for Zika increased across the Americas, scientists developed mathematical models to anticipate further outbreak spread, evaluate possible control measures, and gain insight into outbreak dynamics. These models used a range of data sources including case counts, vector abundance and distribution, population age structure, human mobility, climate information, viral sequence and serological data, and internet ‘big data’ streams. A range of statistical and mathematical models predicted the spread and other epidemic dynamics of ZIKV, as well as the burden of its complications (21–26).

While the WHO PHEIC status was lifted in November 2016 and the neotropical Zika pandemic has waned, the forecasting activities during the pandemic have not been systematically examined, particularly whether the studies were published in a manner and time-frame that was actionable during the Zika pandemic (27). Such an exercise is critical, not only due to the ongoing risk of Zika globally (28), but also to inform modeling efforts for future major epidemics. We therefore undertook a systematic review to identify all published ZIKV prediction and forecasting studies during a time period which encompassed the PHEIC period and the peak and waning phase of the epidemic in the Americas. The first aim of this systematic review was to identify all published models that predicted, forecasted or simulated any ecological or epidemiological phenomenon about the Zika pandemic and describe the predicted phenomena, the range of data sources used and the modeling methods employed. This first aim sought to characterize the methods and data employed to answer key questions during the epidemic and to identify potentially underutilized data or methods. The second aim was to evaluate key scientific characteristics of these studies, including (i) accessibility and timeliness of the publication, (ii) reproducibility of the methods and access to the statistical code and data, and (iii) clarity of the presentation of the prediction results, including uncertainty in prediction estimates. The third aim was to describe the funding structure and major contributing sectors, such as government, industry, non-governmental organizations, or academia, behind these publications.

## Methods

The PRISMA and Cochrane systematic review guidelines were adopted (29). A panel of 12 investigators developed the systematic review protocol including the eligibility criteria and the data abstraction tool. No formal protocol was published for this systematic review.

### Literature search strategy

We conducted a literature review using EMBASE and MEDLINE (PubMed) to identify all potentially eligible studies, which predicted or forecasted phenomenon of the ZIKV pandemic. In MEDLINE we performed a highly sensitive search solely using the term “Zika”. A complementary search in EMBASE used a more specific ontology: “Zika AND (forecasting OR prediction OR model OR modeling OR modelling OR risk OR estimating OR dynamics) NOT mouse”. Both database searches were limited to articles published as of March 1, 2017, and the MEDLINE searching was restricted to those publications released between February 1, 2016 and March 1, 2017. We complemented these database search results with ‘grey literature’, including hand-searching of bibliographies of major Zika epidemiological review articles (17, 30, 31) and contacting experts in the field of Zika modeling to identify any studies which we may have been missed by the above search strategies.

### Screening and eligibility determination

Using a two-reviewer system (with consensus for disagreements and conferral with a 3^rd^ party adjudicator if a consensus was unable to be reached), all articles identified through the above literature search were screened by reviewing the title and abstract to remove all articles that clearly did not meet the eligibility criteria (below). The full text of the remaining articles was reviewed by two reviewers, with a third reviewer if a consensus was not reached by the first two reviewers. Eligibility was based on the following inclusion and exclusion criteria:

### Inclusion criteria

Forecasted, predicted or simulated any epidemiological or ecological phenomenon about the Zika pandemic (including studies regarding previous outbreaks and epidemics, and regions outside the Americas), including but not limited to spatial spread risk, host and ecological range, disease and complication burden, economic impact transmission and other epidemic dynamics. We didn’t require studies to explicitly present a future phenomenon risk, and we included time agnostic estimations of key epidemic parameters and other phenomena.

### Exclusion criteria

- Did not include original analyses (e.g. review articles, perspective pieces, editorials, recommendations, and guidelines)
- Duplicated studies
- Animal and mosquito in-vivo pre-clinical models (e.g mouse, non-human primates)
- *In vitro* studies
- Descriptive epidemiological publications (e.g. describing case positive proportions, total case numbers, descriptive mapping of incidence by geographic information systems)
- Models which only examined causality of ZIKV in Guillain-Barré Syndrome (GBS) or microcephaly (rather than estimating risk or burden, for example)
- Studies which only modeled non-ZIKV arboviruses, unless the central aim of the study was to explicitly forecast or predict ZIKV phenomenon based on the known dynamics of other arboviruses

### Data abstraction, collation and analysis

Data were abstracted from the full texts by 12 reviewers (single-reviewer abstraction) across the domains of (i) objectives and study population, (ii) methodology and reproducibility, (iii) accessibility, timeliness and other bibliometrics of eligible studies, and (iv) author affiliation and funding sources (Table S1). In addition, the availability of preprint manuscripts was assessed using the pre-print search webtool *search.bioPreprint* (32), a server which identifies preprints from arXiv, bioRxiv, F1000Research, PeerJ Preprints, and Wellcome Open Research.

Additionally, we manually searched arXiv and bioRxiv archives to confirm pre-print availability. These pre-print repositories are distinct from the advanced electronic publications made available by most journals after acceptance and peer review. Such ‘grey literature’ review extended beyond the cut-off date for the main literature database searches. A two-reviewer approach was used to ascertain whether eligible studies were made available as pre-print. From the abstracted data, descriptive analyses (medians, IQR, ranges and proportions) and limited hypothesis testing were performed using Stata version 13.0 (StataCorp, College Station, TX, USA).

## Results

Of 2034 studies identified, 73 articles published predominantly from 2016 to 2017 met the inclusion criteria (Fig 1) (20–26, 28, 33–97). The most commonly predicted phenomena were spatial spread (34%), followed by R_0_ (basic reproductive number) or R_E_ (effective reproductive number) (29%), epidemic dynamics (peak size/timing, final size and trajectory) (28%), microcephaly burden (15%), and vector competence and ecology (12%) (Table 1). Most of the geographically resolved predictions were concentrated in the Americas (42%) and Asia-Pacific (21%), while few studies were from Africa (4%). Across 73 studies, the most commonly used data were infection case counts, vector data, and demographic data, followed by climate, meteorological, earth science and transport data (Table 2). Genomic data was not used in any of the studies and few studies used novel real-time internet data streams such as those harnessing open access social media and internet-search engine platforms.

**Figure 1.**
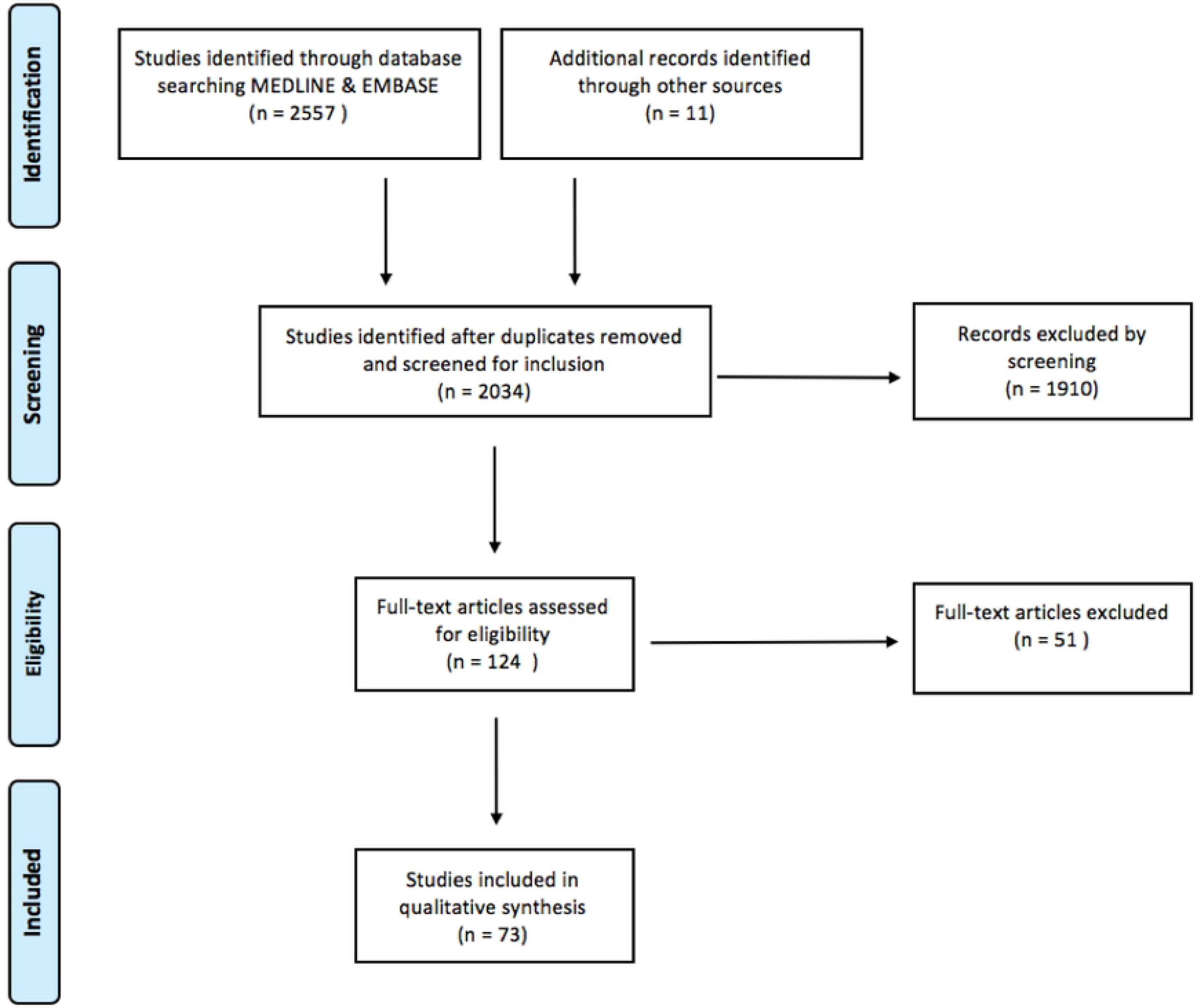
PRISMA flow-chart indicating the number of studies identified, screened and confirmed for eligibility into this systematic review

**Table 1.**
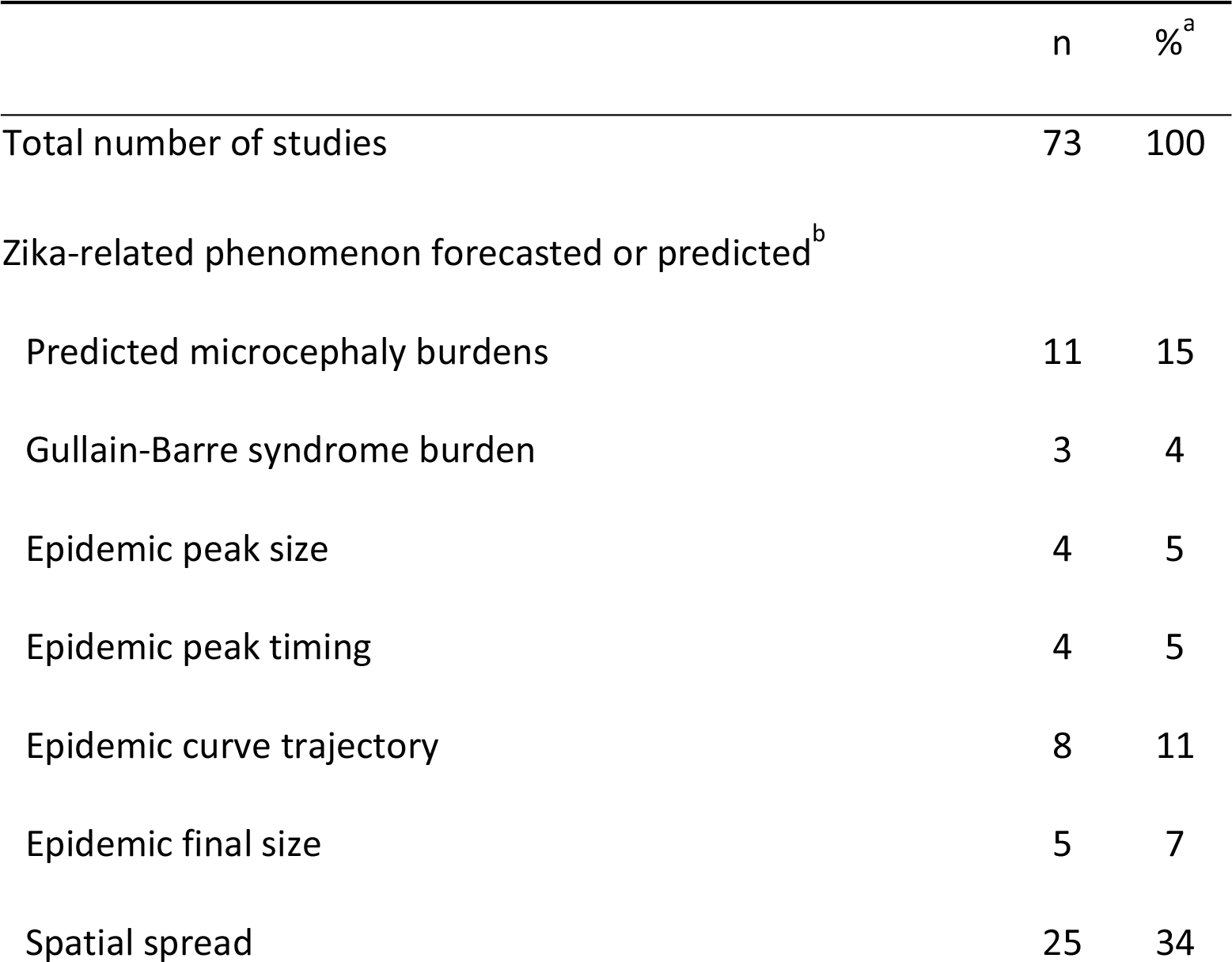
Objectives and study population of eligible studies.

|  | n | %<sup>a</sup> |
| --- | --- | --- |
| Total number of studies | 73 | 100 |
| Zika-related phenomenon forecasted or predicted<sup>b</sup> |  |  |
| Predicted microcephaly burdens | 11 | 15 |
| Gullain-Barre syndrome burden | 3 | 4 |
| Epidemic peak size | 4 | 5 |
| Epidemic peak timing | 4 | 5 |
| Epidemic curve trajectory | 8 | 11 |
| Epidemic final size | 5 | 7 |
| Spatial spread | 25 | 34 |

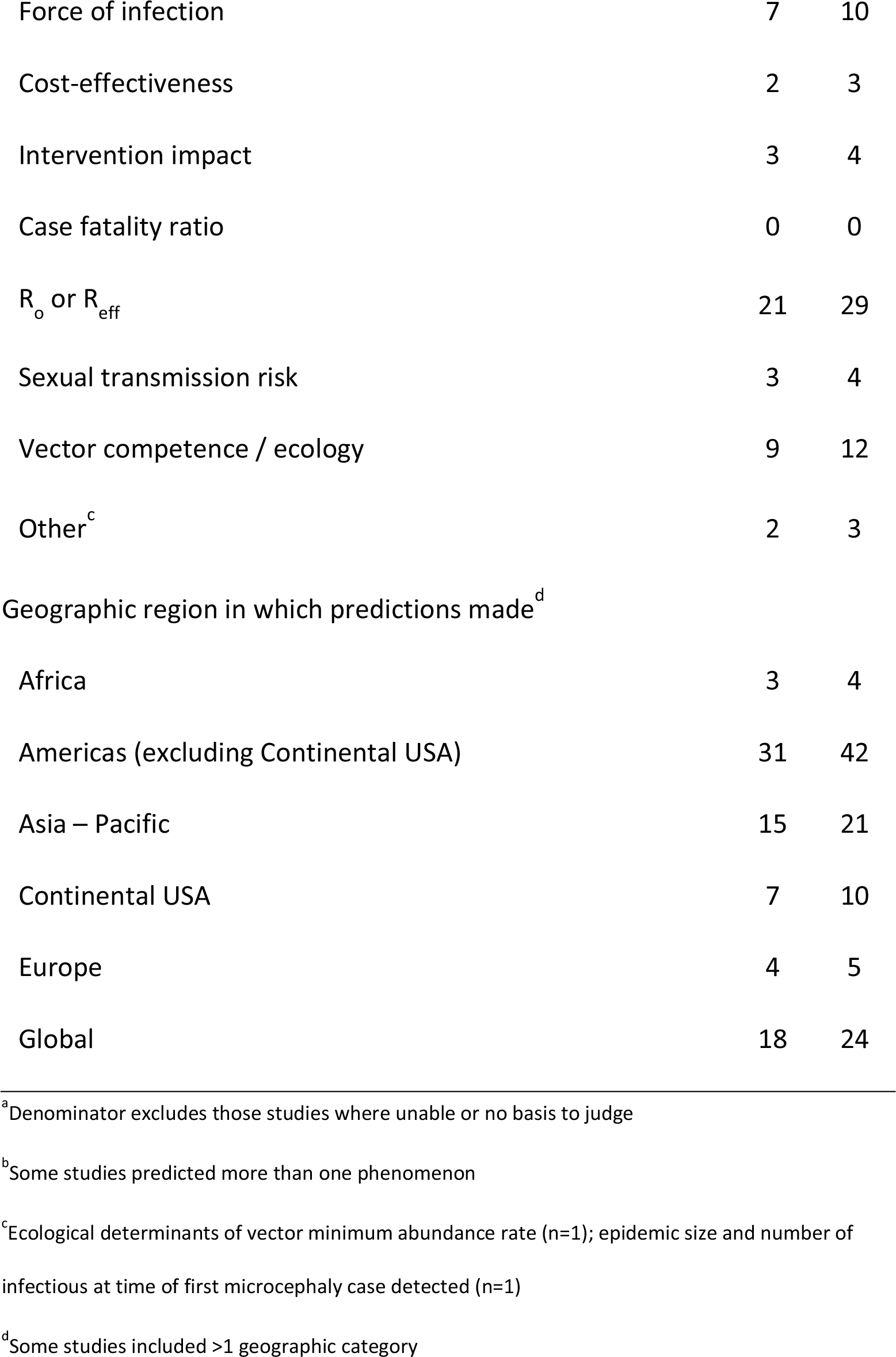

**Table 2.**
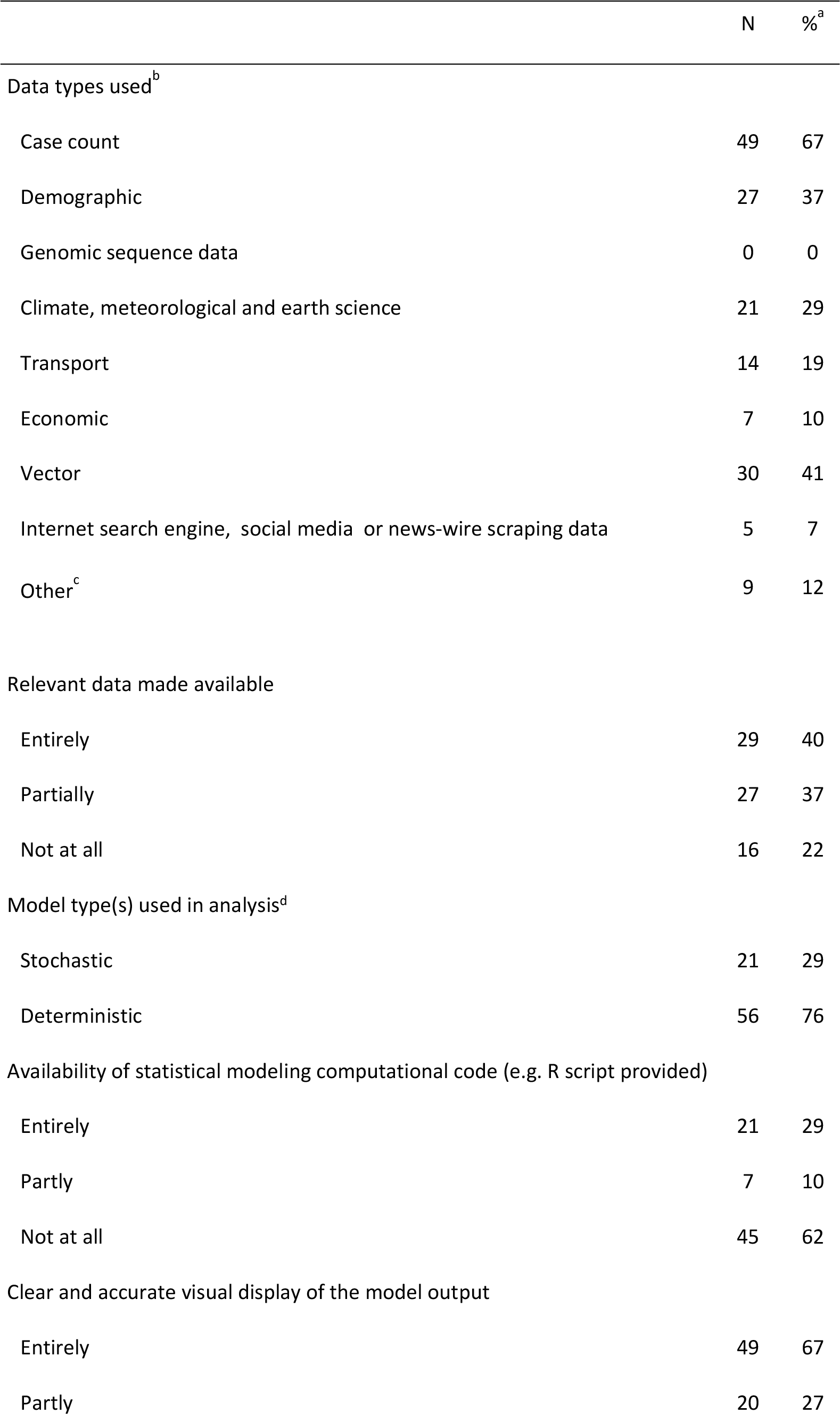
Data sources, methodology and reproducibility of eligible studies.

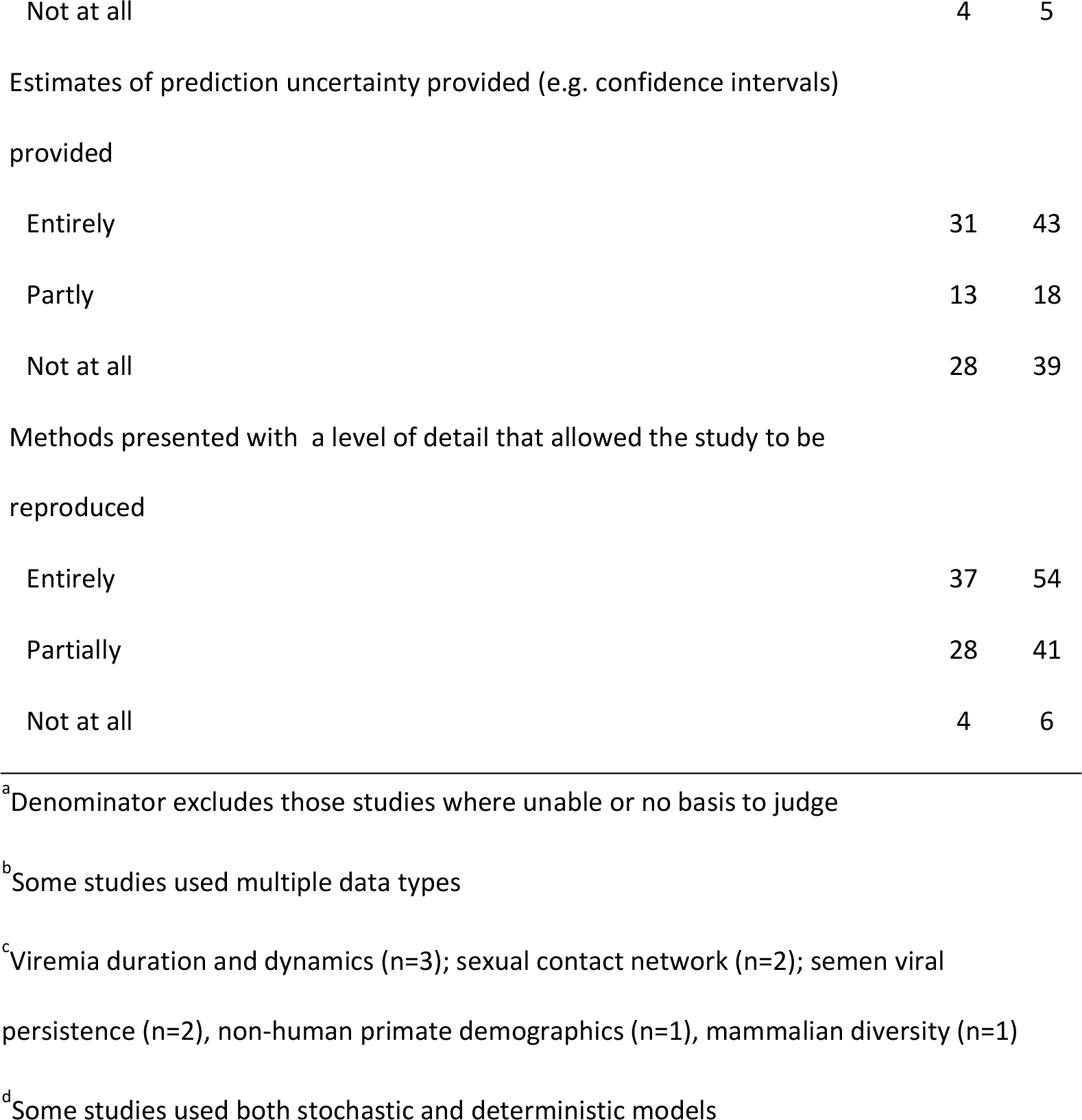

Only 40% of studies made all relevant source data entirely accessible, while more than 20% of the eligible studies did not make any source data available either directly (e.g. an associated data repository) or indirectly (e.g. a citation or web-link) (Table 2). The visual display of model output was at least partly clear and accurate in 95% of the studies. Over a third of the studies did not present estimates of prediction uncertainty. Approximately half of the studies did not entirely present methods with a level of detail to allow reproducibility. Over 60% of the studies did not provide any computational code used for the analyses. We classified more models as deterministic (76%) as opposed to stochastic. It should be emphasized we only ultimately evaluated whether a model was deterministic versus stochastic.

The large majority of published manuscripts were freely accessible (e.g. without a paywall), although 4% were published with paid access only (Table 3). Less than one third of manuscripts were posted on rapid preprint servers (e.g. bioRxiv (98), prior to publication in a peer-reviewed journal. The median time from journal submission to e-journal publication time was 93.5 days, with the maximum time greater than 1 year. This included delays after manuscript acceptance, 25% of the studies had delays of more than 24 days between acceptance and publication (Table 3). Most of the prediction studies were published late in the epidemic, well after the peaks in reported Zika cases (Fig 2, Fig 3). Submitting manuscripts to preprint servers made results available earlier by a median of 119 days (maximum 331 days, IQR 30 – 177 days) (Table 3). This shift led to more results being available close the time of the 2016 South America and Central America epidemic peaks and prior to the epidemic peak in the Caribbean and the 2017 peak in Central America (Fig 2, Fig 3). Comparing the impact factor of journals accepting studies which were posted as preprints (versus the impact factor of those journals accepting studies which were not posted as pre-prints), there was no significant difference (median impact factor 4.37 vs. 4.45 respectively; p = 0.84 by Mann-Whitney U test).

**Table 3.** Accessibility, timeliness and other bibliometrics of eligible studies.

|  | n | %<sup>d</sup> |
| --- | --- | --- |
| Open access<sup>a</sup> | 68 | 96 |
| Pre-print access<sup>b</sup> | 22 | 30 |
|  | median | IQR (range) |
| Journal impact factor | 4.37 | 2.65 - 7.62 (1.48 – 79.26) |
| Submission to acceptance time, days | 83 | 44 - 112 (0 - 256) |
| Acceptance to publication time, days<sup>c</sup> | 15 | 7 - 24 (-255 - 279)<sup>e</sup> |
| Submission to publication time, days | 93.5 | 47 - 141 (1 - 389) |

<sup>a</sup>Includes non-journal open access websites. Open access defined as able to be viewed without any payment or institutional journal license
<sup>b</sup>Biorxiv n = 19, ResearchGate n=1, *Bull WHO* rapid journal pre-acceptance pre-print n = 2
<sup>c</sup>Negative values exist as *Bull WHO* articles published upon receipt (within 24 hrs) and then accepted later
<sup>d</sup>Denominator may vary in cases where these metrics were unable to be determined
<sup>e</sup>Publication time based on electronic journal version where available

**Figure 2.**
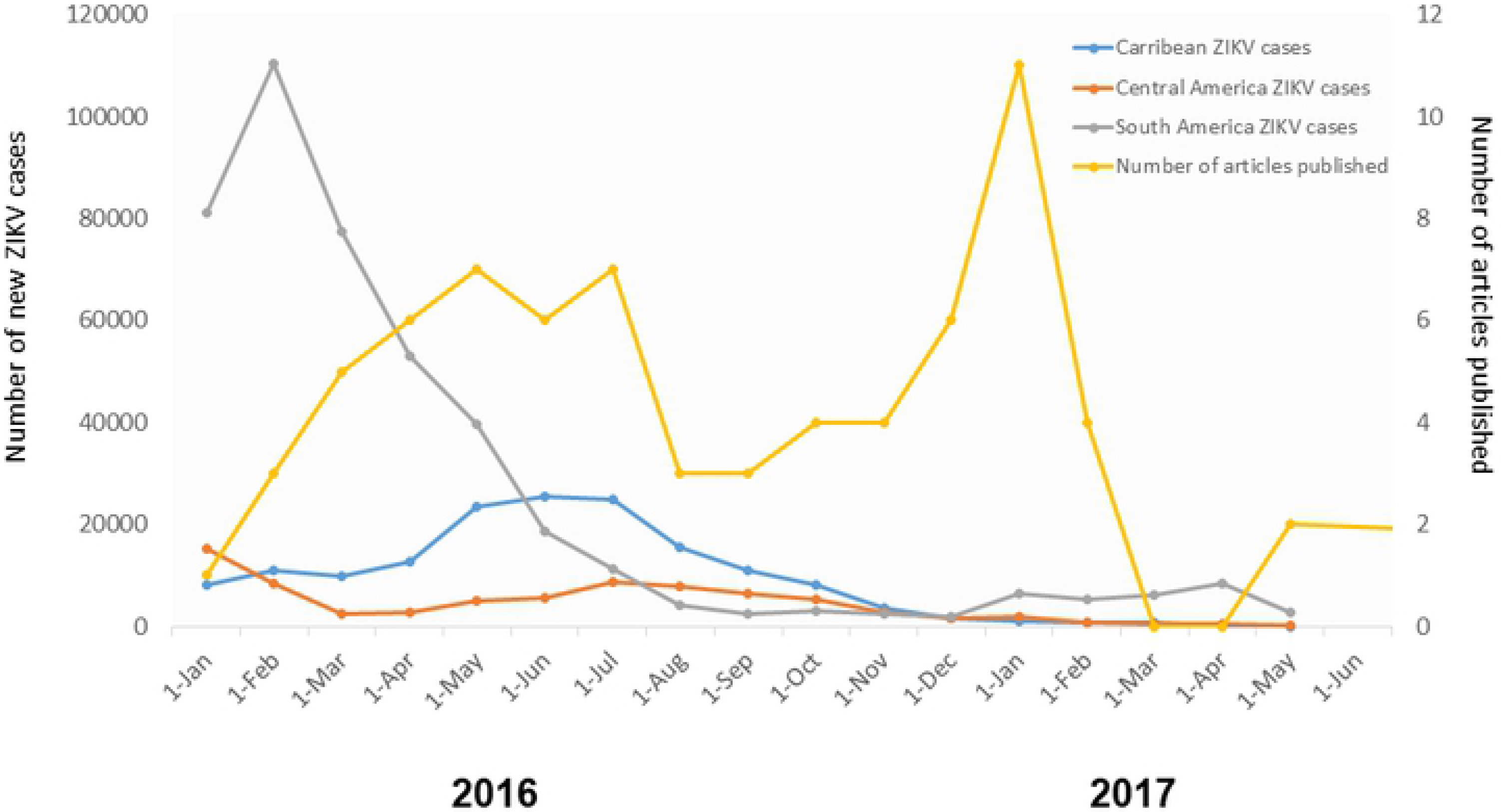
Comparative trends of reported Zika cases in Latin American and publication times of Zika prediction studies. Zika case counts were obtained from https://andersen-lab.com/ with permission

**Figure 3.**
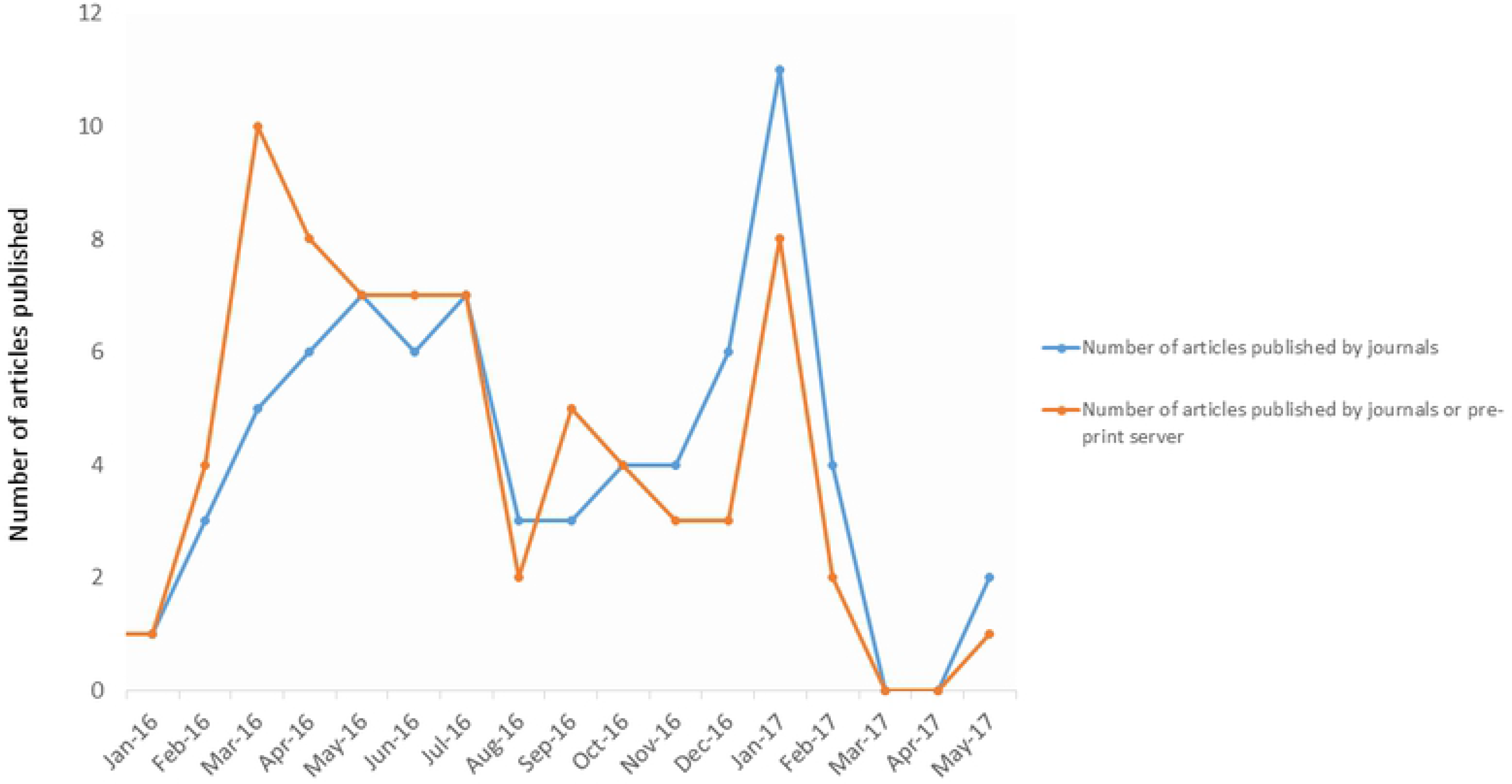
Comparative trends in publication times of ZIKV prediction studies with and without the use of preprints.

Over 90% of the studies included authors with academic affiliations (Table 4). Government affiliated authors participated in a minority of studies, although this may simply reflect “in-house” operational models not being published through journals. Among studies with identifiable funding sources, funding was divided among several sources, though the most common was the United States government, which funded or partially funded 50% of the studies (Table 4). However, many of those studies and other had a variety of funding sources, 85% had at least one non-U.S. government source. Non-governmental organizations were the second most common source, being included in 35% of the studies.

**Table 4.**
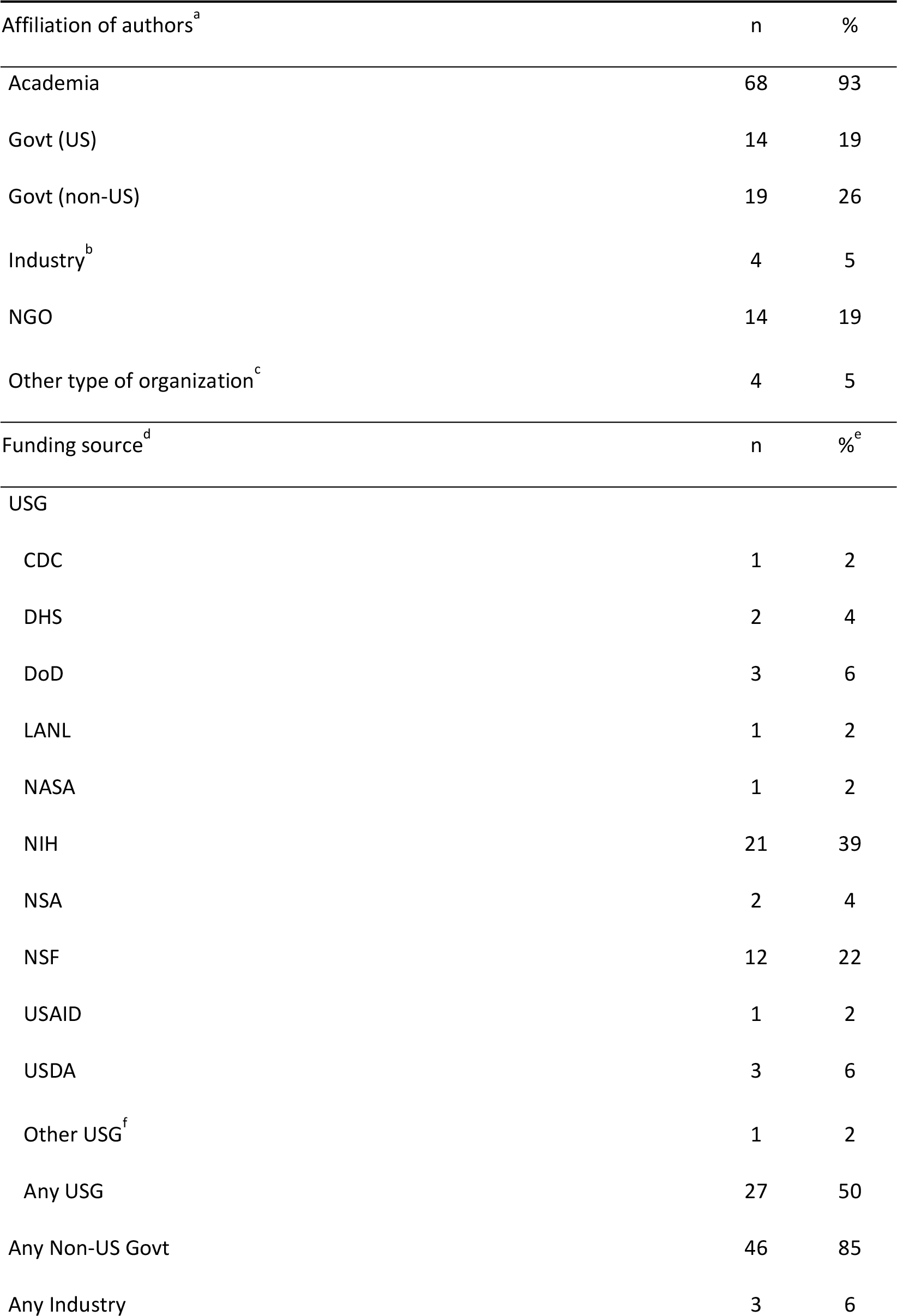
Author affiliation and funding source of eligible studies.

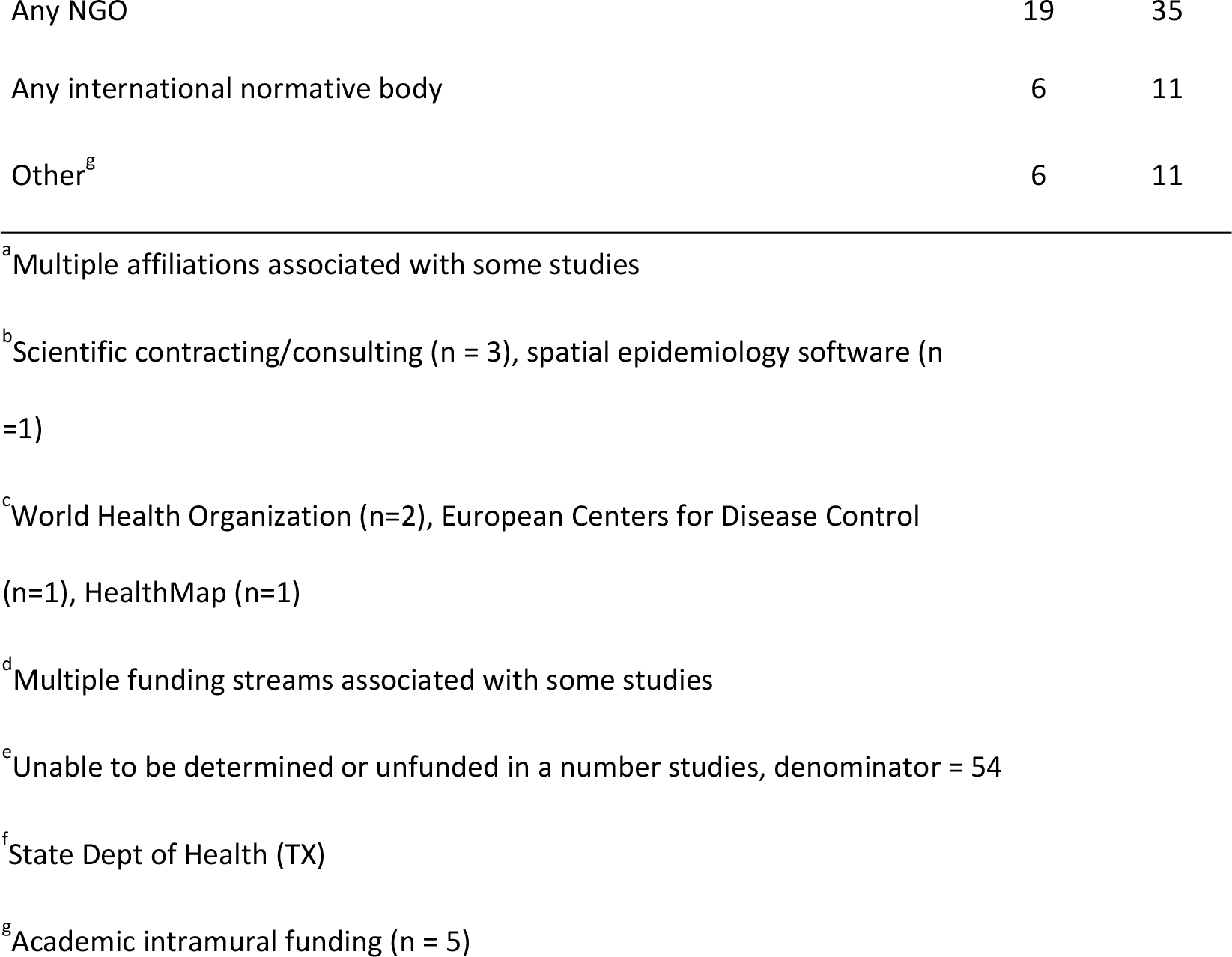

## Discussion

Public health agencies, policy-makers, and other stakeholders are carefully examining the response to Zika. Such ‘lessons-learned’ exercises have been fruitful for prior pandemics and outbreaks, including Ebola, SARS, MERS-CoV, pH1N1, and chikungunya viruses. These exercises have included introspection, analysis, and recommended action with respect to research, public health and policy agendas (99–104). To date, public health ‘lessons-learned’ activities related to the Zika PHEIC have focused on improved ethics preparedness for rapid research during public health emergencies (105), identification of other high-epidemic-risk pathogens with relatively inadequate countermeasure investment (106), expedited approaches to vaccine and other medical countermeasure development (107), rapid data-sharing and material transfer (108–110), and enhancing the role of media communication during epidemics (111).

In contrast to existing reviews on models developed during the ZIKV pandemic, which described specific contributions of modeling (112) or validated analytical assessment of results (113), this systematic review focused on capturing lessons that could improve the functional use of mathematical modeling in support of future infectious disease outbreaks. Extending an approach used by Chretien et al. in their evaluation of Ebola models, we focused on aspects of the studies that likely are particularly relevant to their usefulness during an outbreak (103). This included modeling methods and input data, timeliness and accessibility of the publications, reproducibility (e.g. provision of data and code), and the communication of uncertainty.

Our systematic review identified a large number of Zika models that predicted a wide range of epidemiological and ecological phenomena. The most commonly predicted phenomena were spatial spread, R_0_, epidemic dynamics, microcephaly burden, and vector competence. Notably few of the studies modeled the impact or cost-effectiveness of interventions, sexual transmission risk, or GBS burden. Not surprisingly, the majority of the studies were set in the Americas where most of the cases were reported during the pandemic. Notably one of the global gaps for understanding ZIKV dynamics is Africa, where ZIKV was discovered, is endemic, and poses a risk of future epidemics (114–116).

The leading data types for the examined studies were conventional case counts, vector, demographic, climate and transport data. This finding reflects not only the availability but also the importance of such data. Case count data in particular are often hard to access but critical to many modeling approaches. Rapid sharing of case count data during international public health emergencies, as well as open, curated, rapidly accessible baseline demographic, human mobility, climate, and environmental datasets are essential to quickly leverage modeling and forecasting efforts (109). Our review also identified several relatively underused data streams. First, socioeconomic and behavioral data were conspicuously absent. The lack of behavioral components in these models is concerning given the importance of these factors on disease dynamics. Second, real-time internet-based data-streams, such social-media and internet search-engine data, were used in a minority of ZIKV prediction studies identified in this systematic review. The limited use of internet ‘big data’ in the models suggests that either these data are of lower value for epidemic forecasting or that methods have yet to be developed to efficiently extract important information from them. Such data streams may be more commonly used in forecasting in the future as their strengths and weakness become clearer (117).

Genomic data were absent from these published models. During the pandemic, sequencing platforms were employed to generate data critical to diagnostic and countermeasure development (118), but our systematic review revealed that these data were not incorporated into prediction frameworks during the first year of ZIKV pandemic. This may reflect that early molecular epidemiology studies aimed to reconstruct the invasion and evolution of ZIKV rather than forecasting future changes (119, 120). Some phylodynamic studies were published after the time period of the systematic review, with interesting results highlighting the possibility for phylogenetic data to provide unique insight into epidemic dynamics and possibly forecasting (120–122). The relative delay of these studies (relative to other to those using other data sources) echoes a similar time lag of phylogenetic studies during the 2015 Ebola epidemic (103). The lack of phylogenomic studies captured by this review also suggests that substantial bottlenecks still exist in using these data sources in epidemic response, despite advances in mobile near “real-time” sequencing technologies (118). In the future, as new methods are developed, and genomic data become more readily available, the use of these data will likely become more common in prospective forecasting frameworks.

Our systematic review did not delve deeply into modeling approaches, but did identify a preponderance of deterministic as opposed to stochastic models. Both categories of models have pros and cons and their use is often informed by the specific question being addressed, in addition to data availability (123). Deterministic models may generally be easier to produce, but they do have limitations for intrinsically stochastic processes like epidemics, such as underestimating uncertainty (124). Uncertainty is particularly important in this context where uncertainties are generated by the epidemic itself, data collection, and analytical approaches. Moreover, forecasts are ideally used to inform the mobilization of resources to save lives, a context in which clearly characterizing uncertainties is paramount. This is also a clear area for improvement in model output reporting; only 43% of studies completely reporting uncertainty.

Our review also provided a unique evaluation of the more functional aspects of published predictions and forecasts. We determined that the visual clarity of model output was high but indicate room for improvement in publishing datasets used for model fitting and validation, sharing computational code for others to potentially rapidly implement the model, presenting estimates of prediction uncertainty, and methodological detail to allow the study to be reproduced. The variable quality in sharing model code and methodological detail shown here does suggest that epidemic model reporting consensus guidelines, which establish a minimum standard for the reporting of epidemic modeling, may be valuable. A recent review of the modeling efforts for the Ebola epidemic also called for standardization of modeling practice (103). Many other fields of biomedical research have established reporting guidelines to improve research quality and implementation (125–128).While reporting guidelines have been proposed for population health model on a broader scale (129), none have been established for epidemics.

This review also indicated that a majority of studies (60%) did not completely disclose the data they used. To the extent permissible with ethical and privacy constraints, publishing the aggregated data used to fit and validate models is critical. Not only would sharing data support full reproducibility, but sharing would also enable other researchers to use data in their own complementary modeling efforts. Modelers could therefore help answer calls for increased data sharing during public health emergencies (103, 109, 130). Exploring how data can be shared more openly and quickly during a public health emergency would be useful, as this remains a challenge.

Many studies identified in this review were published on a time-scale that was relevant to the Zika response. However, a large number of predictions were published well after the epidemic peaks, limiting their ability to inform the response. Nonetheless, those studies may well be used to inform other preparedness activities and contributed to the general knowledge of the biology, epidemiology and/or ecology of ZIKV. Further, results may have been informally shared with public health officials or other relevant decision makers prior to publication. Similar delays to publication have also been noted in an analysis of modeling efforts during the 2015 Ebola epidemic, which noted a median publication lag of around three months [103].

We identified two modifiable bottlenecks in the dissemination of results. First, delays from acceptance to journal publication were generally minimal (median 15 days), but a quarter of the evaluated studies had greater than 24 days delay from journal acceptance to publication. Immediate posting of accepted papers, as practiced by many journals, could cut this time down substantially. Second, we found that only 30% of studies were made available as preprints prior to peer review despite endorsements of preprints by major public agencies, funders, and journals. Those posted were available a median of 119 days prior to peer-reviewed publication. An analysis of preprints for all Zika publications over a similar time period found similar publication delays but much lower overall preprint use compared to the studies analyzed here (3.4% versus 30%) (131). This greater adoption may indicate a changing preprint culture which was also reflected by our finding that preprint posting did not have a demonstrable effect on the impact factor of the journal in which the study was published, and we suggest that pre-prints be more frequently used in future public health emergencies, echoing other similar recent arguments (131).

Our review also provided a unique analysis of the funding sources and author affiliations of the published ZIKV prediction and forecast efforts across the ZIKV pandemic. These results indicated a range of stakeholders. We note that while academia contributed to the greatest volume of published studies, our search strategy would not have captured in-house models developed by US federal agencies or other unpublished models which may have provided direct operational support.

This systematic review has three important weaknesses. First, due to scale, a completely independent two-reviewer system was not used for abstracting most of the data and for evaluation of aspects such as reproducibility. Second, we did not formally search for preprint manuscripts as part of the literature searching phase of the systematic review, only assessing whether eligible manuscripts had corresponding preprints. We may have therefore missed important research that had been posted but not yet peer-reviewed. Lastly, we had to restrict the time frame for publications to consider in the review. This restriction again led to missing studies, some of which may have already been published but not yet posted in EMBASE or MEDLINE.

Overall, the review identified several areas of improvement such as providing data and code, developing reporting standards, posting preprints, and communicating uncertainty. Addressing these areas can improve our understanding of Zika and other outbreaks and ensure that forecasts can inform preparedness and response to future outbreaks, epidemics and pandemics.

## Acknowledgements

The authors wish to thank members of the Pandemic Prediction and Forecasting Science and Technology Working Group for their helpful feedback. The authors also acknowledge the valuable curated Zika case count data used with permission by the Andersen lab (https://andersen-lab.com/).

## Disclaimer

The views expressed in this article are those of the authors and do not necessarily reflect the official policy or position of the Department of the Army, Department of Defense, USUHS, nor the U.S. Government. Several of the authors are US Government Employees. This work was prepared as part of their official duties. Title 17 U.S.C. § 105 provides that ‘Copyright protection under this title is not available for any work of the United States Government.’ Title 17 U.S.C. §101 defines a U.S. Government work as a work prepared by a military service member or employee of the U.S. Government as part of that person’s official duties.

The contents of this publication are the sole responsibility of the authors and do not necessarily reflect the views, assertions, opinions or policies of the Uniformed Services University, the Department of Defense, the Centers for Disease Control & Prevention, or US Environmental Protection Agency. Mention of trade names, commercial products, or organizations does not imply endorsement by the U.S. Government. More than one author is an employee of the U.S. Government and as such under the provisions of 17 U.S.C. 105, copyright protection is not available for this work

## Supporting Information Legends

**Table S1.** Data abstraction and study evaluation tool used by reviewers

